# Post-transcriptional control contributes to hypoxia-induced tumorigenic phenotypes in macrophages

**DOI:** 10.1101/2023.10.09.561558

**Authors:** Edward M.C. Courvan, Roy R. Parker

## Abstract

Macrophages are effector immune cells that experience substantial changes to oxygenation when transiting through tissues, especially when entering tumors or infected wounds. How the transition to hypoxia alters gene expression and macrophage effector function remains poorly understood, especially at the post-transcriptional level. Here we use TimeLapse-seq to measure how hypoxia modifies inflammatory activation of primary macrophages. Nucleoside recoding sequencing allowed us to derive steady-state transcript levels, degradation rates, and transcriptional synthesis rates from the same dataset. We find that inflammatory activation of macrophages is altered by hypoxia due to increased mRNA decay. Destabilized transcripts encode for proteins associated with phagocytosis and proteolysis of the extracellular matrix. Consistent with differential gene expression, we observed hypoxia alters the specificity of macrophage phagocytosis and reduces their invasion of extracellular matrix. Hypoxic gene expression bears similarity to tumorigenic macrophages in solid tumor biopsies suggesting post-transcriptional control contributes to the macrophage transition from tumoricidal to tumorigenic.

## Introduction

Macrophages are a pillar of the mammalian innate immune system. They have a canonical role in phagocytosing damaged cells and pathogens, but also coordinate the response of other immune cells in an effector role^1^. Macrophages sense the physical and chemical environment for stimuli such as cytokines, complement system ligands, and oxygen levels so that they may integrate them into the appropriate response. Though biochemical cues (i.e. cytokines) have been well studied for their effects on macrophages, we know less about how environmental factors like hypoxia lead to specific gene expression patterns in macrophages and how that affects downstream consequences for surrounding tissues^1^. Such responses are essential to proper wound healing and infection resolution but can backfire in solid tumors where macrophages frequently adopt a regenerative phenotype advantageous to tumor growth^2, 3^.

*In vitro* studies have historically used a two-state model of activation whereby macrophages are polarized to be inflammatory (M1) or regenerative (M2)^4, 5^. It is now understood that macrophage polarization is a far more complex landscape of interconnected pathways, and that specific stimuli must be examined in light of how they modify classical gene expression patterns^6^. Recent work has shown that, in human tumor samples, the M1/M2 axis breaks down entirely and might be better replaced by other biomarkers for macrophage tumoricidal or tumorigenic phenotype^7^. Contemporary sequencing methods have the power to reveal how mixed stimuli converge to determine the macrophage phenotypes observed in real tissue and more realistically refine the classical two-state paradigm.

Macrophages encounter hypoxia in tissues and solid tumors. Indeed, hypoxia in tumors is thought to disable some macrophage anti-tumorigenic properties and confer resistance to treatment^8–10^. A great deal of effort has been invested in understanding transcriptional regulation in hypoxia, but we are only beginning to understand how post-transcriptional regulatory mechanisms affect mRNA function and stability in hypoxia^11–13^. Studying post-transcriptional regulation in macrophages and other immune cells is critical because mRNA decay and stabilization can exert significant effects on transcript levels over a much shorter timespan than transcription alone^14, 15^. If we are to understand how macrophages respond to changes in their environment, we must understand the processes that govern transcript lifetime.

Here we present data showing that post-transcriptional regulation leads to widespread transcript destabilization in hypoxic inflammatory macrophages. We deploy TimeLapse-seq to measure transcriptome-wide degradation rates (k_deg_) in primary human macrophages that are subjected to hypoxia with simultaneous induction with lipopolysaccharide (LPS) and interferon gamma (IFNγ). We find that hypoxia modifies the expression of inflammatory signals and downregulates transcripts related to motility, tissue invasion, and phagocytosis via increased mRNA decay. Altered mRNA levels lead to measurable differences in phagocytic specificity and extracellular matrix (ECM) digestion within 24 hours of hypoxic onset. Comparing our results to published single cell RNA-seq, we find that hypoxia drives inflammatory macrophages towards a regenerative state that recapitulates a subset of tumorigenic macrophage gene expression signatures.

## Results

### TimeLapse-seq and BakR capture transcriptional kinetics of inflammatory activation

We chose to analyze the effects of hypoxia on the inflammatory activation of macrophages (M1 in the canonical framework) using the TimeLapse-seq modality^16^. Briefly, TimeLapse-seq includes a 4-thiouridine (s^4^U) feed at the end of a time course, which is incorporated into newly transcribed RNA. After harvesting and RNA isolation, the s^4^U is treated by an oxidant that recodes the hydrogen base pairing at the Watson-Crick face to that of a cytidine. After sequencing, U to C mutations can be scored to determine which reads are from an RNA transcribed before or after the s^4^U feed. The proportion of computationally sorted reads then allows for statistical inference of the mRNA degradation rate for each transcript. Assuming the sample is near steady-state, the synthesis rate (k_syn_) may be inferred as well.

We verified that primary human macrophages uptake s^4^U added to media using a fluorescence conjugation assay and confirmed that mutations are detected in samples that are fed s^4^U but not an unfed control (Figure S1A&B). We find that unfed cells have a transcriptome-wide uridine mutation rate of less than 0.08%, while macrophages fed with 500 μM s^4^U for two hours generate a mutation rate of 1.63% which is more than three standard deviations away from our unfed control – ample signal for determining transcript degradation rates (Figure S1C).

Our experiment compares three conditions: 1) resting macrophages in atmospheric oxygen, 2) macrophages activated with LPS and IFNγ in atmospheric oxygen, and 3) macrophages activated with LPS and IFNγ in hypoxic conditions. Atmospheric oxygen at the elevation of our lab is approximately 120 mmHg, equivalent to 17% at sea level. Our hypoxic incubator was calibrated to 7 mmHg oxygen, 1% at sea level. We chose these conditions because 1% is a physiological level of hypoxia where oxidative phosphorylation may still proceed and is found in many tissues and tumors^17, 18^. Each condition was harvested in triplicate at twenty-four hours after treatment. All samples were fed with 500 μM s^4^U for the last two hours of incubation (Figure 1A).

**Figure 1.**
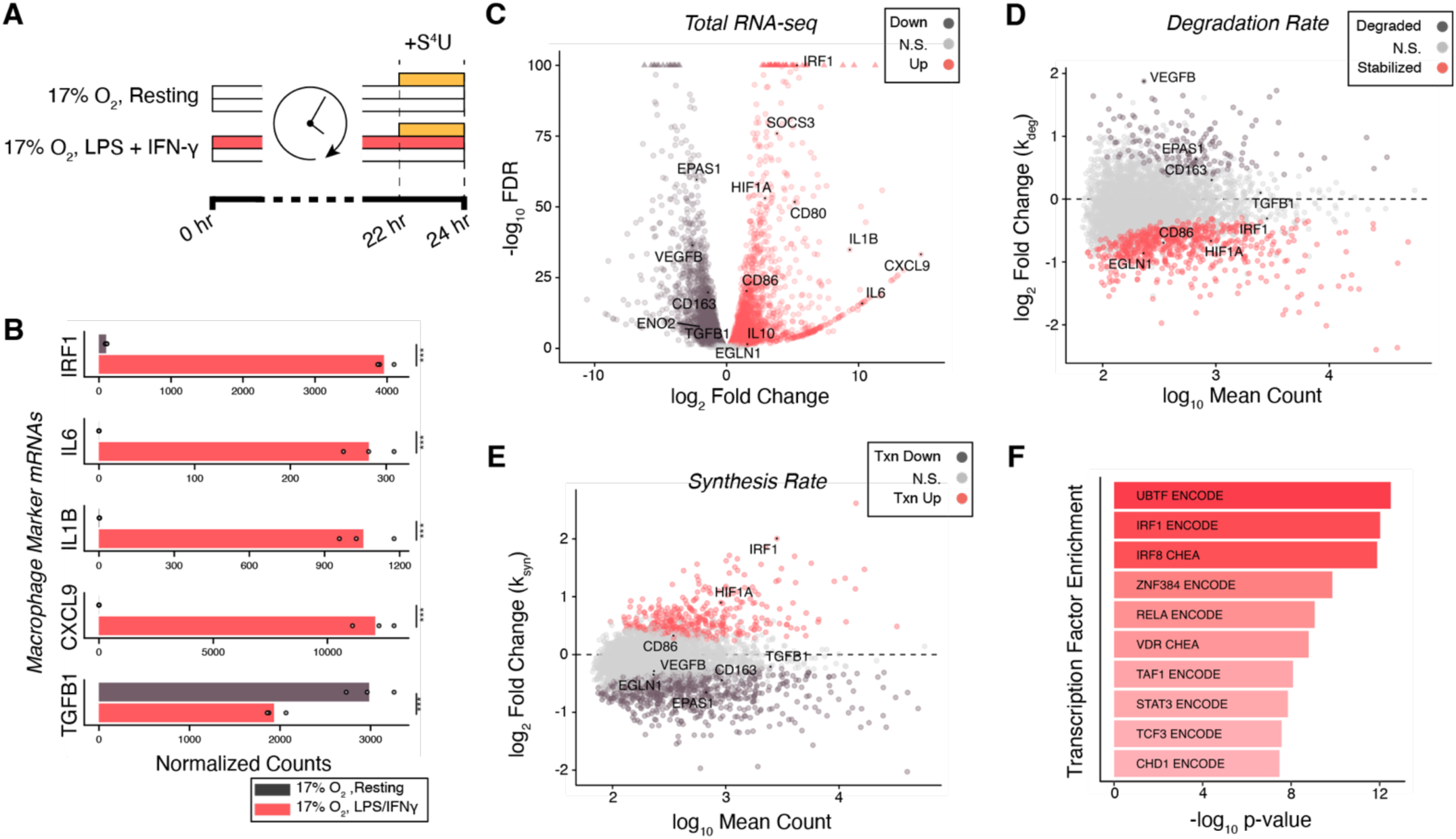
TimeLapse-Seq reveals the mechanism of gene expression in macrophage activation. A) Experimental design for TimeLapse-seq. Macrophages were incubated for 24 hours where the activated samples were either left untreated or treated with LPS and IFNγ (red) in room oxygen of 17% O_2_. s^4^U labeling was applied in the last two hours (yellow). B) Normalized read counts for specific genes related to macrophage activation where points are individual replicates and bars represent the mean of those replicates. FDR threshold given as: *** FDR < 0.001. C) Total RNA expression verses false discovery rate determined by DESeq2 between resting and inflammatory activation. D) Abundance versus differential degradation rate (k_deg_) between resting and inflammatory activation. Significance threshold set to FDR < 0.05. E) Abundance versus differential synthesis rate (k_syn_) between resting and inflammatory activation. Significance threshold set to FDR < 0.05. F) Top ten transcription factors with enriched promotor binding sites for transcripts with increased synthesis. Enrichment is drawn from the consensus database of ENCODE and ChEA transcription factors.

We analyzed our data for changes in steady-state RNA levels using DESeq2 and changes in transcript degradation rate using bakR^19, 20^. DESeq2 is a widely accepted method for differential expression analysis in RNA-seq methods whereas bakR is a package for analyzing nucleoside recoding data using Bayesian Hierarchical modeling. Briefly, bakR infers the degradation rate of a transcript from the fraction of mutated uridines in each mapped read. Combining degradation rates with differential expression determined by DESeq2, one can compute transcript synthesis rates and therefore determine whether synthesis or stabilization drives changes in expression.

We first sought to benchmark our TimeLapse-seq against prior findings in macrophages stimulated by LPS and IFGψ in normoxia. We observed that LPS and IFNγ produce the expected inflammatory response comparing resting normoxic macrophages to activated normoxic macrophages. Specifically, we selected a panel of canonical markers of the LPS/IFNγ response to gauge whether macrophages were activating as expected^6^. Transcription factor IRF1 is strongly induced, as are the inflammatory cytokines CXCL9, IL6 and IL1B (Figure 1B). Transcripts for surface markers of inflammatory macrophages, CD80 and CD86 were also upregulated while CD163, a marker of regenerative macrophages, is downregulated (Figure S1D). TGFB1 and IL10, cytokines associated with a regenerative phenotype, are respectively downregulated or barely detected in normoxic activated macrophages (Figure 1B&S1D). Based on these markers, we concluded that our dataset indeed measures transcriptome changes during inflammatory activation of macrophages in response to LPS and IFNγ in normoxia.

In addition to canonical markers, the macrophage transcriptome changes profoundly upon induction with LPS and IFNγ in normoxia (Figure 1C). We found over 6000 differentially expressed genes (False discovery rate (FDR) < 0.05) where log2 fold-change ranges from -11 to 15 (Figure 1C). Our bakR analysis, independent of differential expression, identifies 746 transcripts with significantly altered degradation rates (FDR < 0.05) ranging from k_deg_ log_2_(fold-change) of -2.4 to 1.9 (Figure 1D). IFNγ can inhibit regenerative pathways like angiogenesis, and the angiogenic factor VEGFB is accordingly downregulated^21^. From our differential degradation data, we can conclude that this downregulation of VEGFB is driven by increased degradation of its mRNA and infer that LPS and IFNγ downregulate regenerative processes like angiogenesis.

By combining differential steady-state RNA transcript levels with changes in degradation rate, we can infer the differential synthesis of each gene (Figure 1E). We measure 1141 transcripts to be significantly differentially transcribed upon inflammatory activation (FDR < 0.05). Notable amongst these is the interferon responsive transcription factor IRF1. The remaining genes of the 6000 for which we measure differential expression were either filtered out of the analysis by bakR, which requires a minimum of 50 reads per experimental condition to estimate decay and synthesis rates, or did not pass our statistical thresholds. Many of the genes with the highest differential expression were not detected in resting macrophages (i.e., IL6, IL1B, CXCL9). Taken in sum, we observe that inflammatory activation is dominated by new transcriptional activity with post-transcriptional dynamics impacting a subset of differentially expressed genes.

To validate our parsing of differential expression into differential degradation and synthesis we used an enrichment analysis strategy to identify transcription factors driving the inflammatory response. We compared genes that were synthesized at significantly increased rates (FDR < 0.05) against the ENCODE and CHEA databases using EnrichR and found that these changes are driven by IRF1, IRF8, and RELA, known drivers of inflammatory transcription (Figure 1F)^22, 23^. Genes with reduced new synthesis enrich for a nearly non-overlapping set of transcription factors suggesting our differential synthesis analysis reflects specific transcriptional reprogramming (Figure S1E).

We conclude that macrophages induced with LPS and IFNγ change their gene program through both transcriptional and post-transcriptional mechanisms that are effectively captured by TimeLapse-seq and bakR.

### Macrophages undergo inflammatory activation in hypoxia

Hypoxia has been reported to regulate macrophage inflammation,^9, 24^ so we compared normoxic resting macrophages to macrophages activated by LPS and IFNγ in hypoxia (Figure 2A). We find that the same marker genes of macrophage inflammation, IRF1, CXCL9, etc., are similarly induced in hypoxia (Figure 2B&S2A).

**Figure 2.**
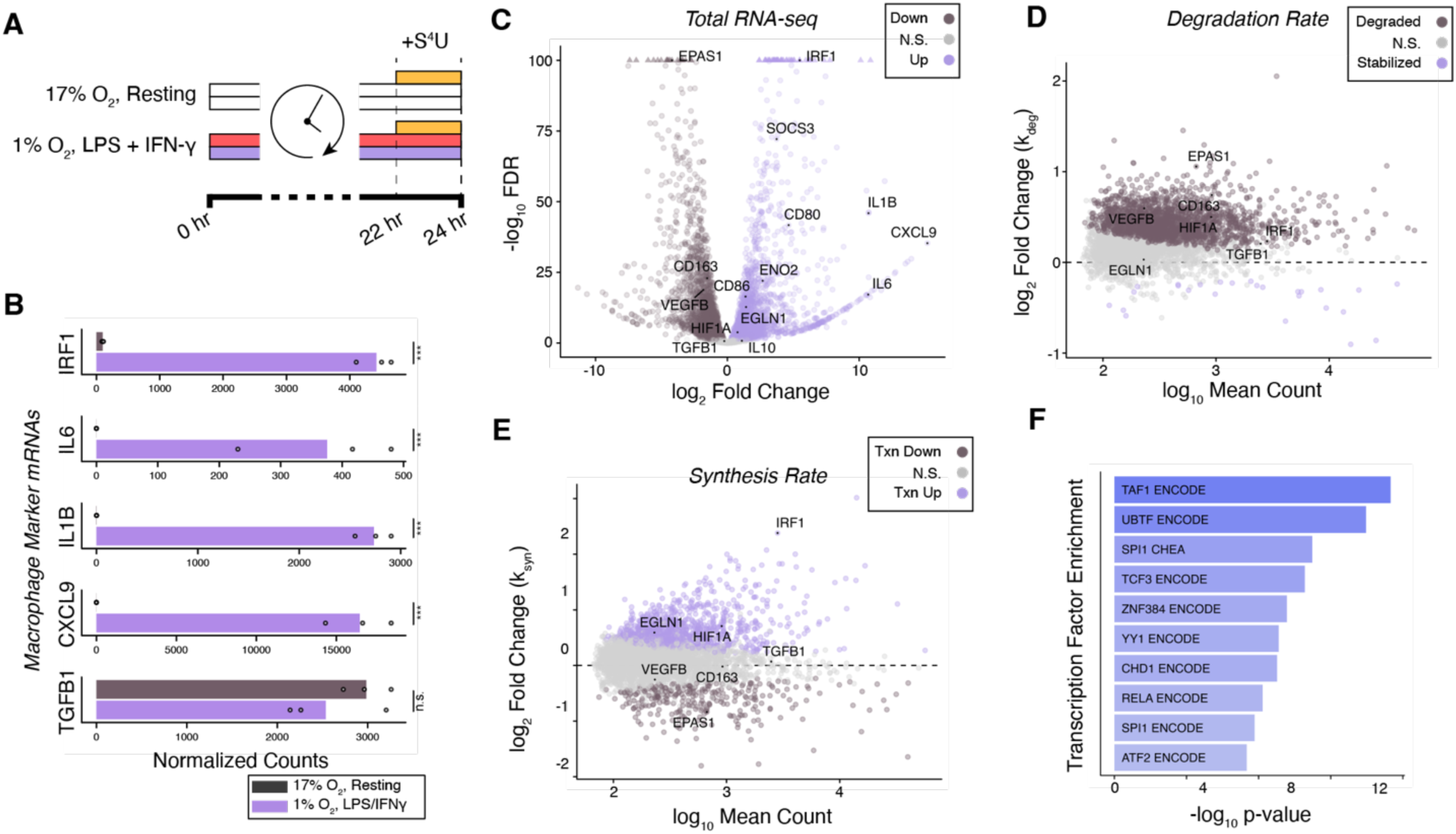
Macrophages are activated in hypoxia but undergo increased RNA decay. A) Schematic of comparison between resting macrophages in room oxygen of 17% O_2_ and macrophages treated with LPS and IFNγ (red) and moved to a hypoxic chamber at 1% O_2_ (purple) for 24 hours. s^4^U labeling was applied in the last two hours (yellow). B) Normalized read counts for specific genes related to macrophage activation where points are individual replicates and bars represent the mean of those replicates. FDR thresholds given as: *** FDR < 0.001 and n.s. as not significant with FDR > 0.05. C) Total RNA expression verses false discovery rate determined by DESeq2 between resting and inflammatory activation. D) Abundance versus differential degradation rate (k_deg_) between resting and inflammatory activation. Significance threshold set to FDR < 0.05. E) Abundance versus differential synthesis rate (k_syn_) between resting and inflammatory activation. Significance threshold set to FDR < 0.05. F) Top ten transcription factors with enriched promotor binding sites for transcripts with increased synthesis. Enrichment is drawn from the consensus database of ENCODE and ChEA transcription factors.

Hallmarks of the hypoxic response are apparent in our comparison of normoxic resting macrophages and hypoxic LPS and IFNγ activated macrophages. The canonical hypoxia inducible genes HIF1A and EPAS1 (also known as HIF2A) are moderately downregulated while their targets EGLN1 and ENO2 are upregulated (Figure 2C). EGLN1 encodes a prolyl hydroxylase that targets HIF1A as part of an autoinhibitory loop regulating the hypoxic response, and ENO2, encoding enolase, reflects a shift towards glycolytic metabolism^25–27^.

Strikingly, hypoxia causes many mRNAs to become unstable (Figure 2D). We found that more than 1800 genes were significantly destabilized (FDR < 0.05) while only 26 are stabilized.

Differential synthesis results were largely similar in hypoxia when compared to activation in normoxia, evidenced by the induction of IRF1 (Figure 2E). However, the array of active transcription factors is altered, with YY1 and ATF2, both stress responsive transcription factors, now appearing in the top ten most enriched transcription factors under hypoxic conditions (Figure 2F)^28, 29^.

### Coding transcripts experience widespread RNA decay in hypoxia

To isolate the effect of hypoxia on macrophage activation, we directly compared activated normoxic macrophages to activated hypoxic macrophages (Figure 3A). This comparison showed that inflammatory cytokines IL1B and CXCL9 are even more induced in hypoxia than in normoxia (Figure 3B). Differential expression analysis shows that VEGFB and CXCR4 are upregulated as expected in hypoxia, while PLAU and several integrins are among those downregulated (Figure 3C)^10^. PLAU encodes the urokinase plasminogen activator, which is a major regulator of macrophage adhesion and tissue invasion^30^. Again, EGLN1 and EGLN3, genes encoding PHD proteins which are part of a HIF1a autoinhibitory loop, are highly expressed^27^.

**Figure 3.**
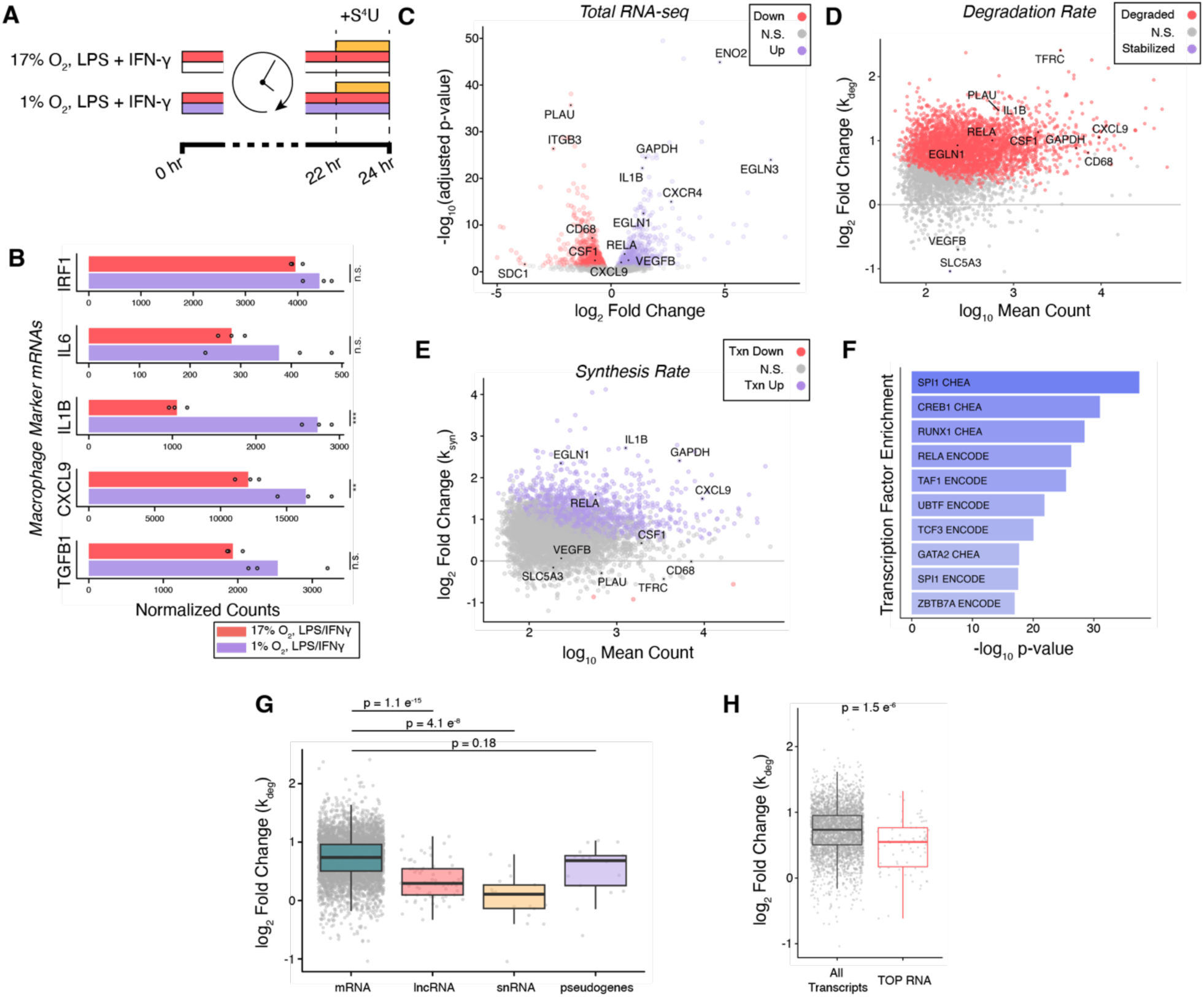
Hypoxia induces widespread degradation of mRNA transcripts. A) Schematic of comparison between macrophages treated with LPS and IFNγ (red) in room oxygen of 17% O_2_ and macrophages treated with LPS and IFNγ and moved to a hypoxic chamber at 1% O_2_ (purple) for 24 hours. s^4^U labeling was applied in the last two hours (yellow). B) Normalized read counts for specific genes related to macrophage activation where points are individual replicates and bars represent the mean of those replicates. FDR thresholds given as: ** FDR < 0.01, *** FDR < 0.001, and n.s. as not significant with FDR > 0.05. C) Total RNA expression verses false discovery rate determined by DESeq2 between resting and inflammatory activation. D) Abundance versus differential degradation rate (k_deg_) between resting and inflammatory activation. Significance threshold set to FDR < 0.05. E) Abundance versus differential synthesis rate (k_syn_) between resting and inflammatory activation. Significance threshold set to FDR < 0.05. F) Top ten transcription factors with enriched promotor binding sites for transcripts with increased synthesis. Enrichment is drawn from the consensus database of ENCODE and ChEA transcription factors. G) Transcripts grouped by type compared to differential stability (k_deg_). Kruskal-Wallis rank sum test p = 1.9 e^-21^. P values reported in plot derived from Dunn’s test. H) Differential degradation of TOP RNA compared to all other transcripts. P value derived from Mann-Whitney rank sum test.

Comparing degradation rates of normoxic to hypoxic inflammatory activation, we again observe widespread mRNA decay. We find that over 3000 out of 5300 transcripts exhibit increased degradation (FDR < 0.05) during inflammatory activation in hypoxia compared to the same activation in normoxia (Figure 3D).

Global effects in the transcriptome can cause DESeq2 and other differential expression algorithms to miscalculate the magnitude of differential expression because it assumes that most of the transcriptome is unchanged during a perturbation^31^. We examined two lncRNAs, MALAT1 and XIST, both of which are expressed at high levels and should be relatively unchanged by hypoxia. As expected, their expression levels remain flat, even though they turn over more rapidly (Figure S3A). This argues that DESeq2 had not introduced a systematic skew to differential expression in hypoxia.

The destabilization of so many transcripts prompted us to ask whether any distinguishing feature correlated with that destabilization. We separated transcripts into deciles according to the k_deg_ measured in normoxic inflammation and observe no skew towards more destabilization based on k_deg_ of the transcript in normoxic inflammation (Figure S3B). Thus, both stable and unstable mRNAs show faster degradation rates in hypoxic conditions. Separating transcripts by class, we find that mRNAs are substantially more degraded than non-coding RNAs (Figure 3G). We interpreted this observation to suggest that hypoxia specifically causes increased decay of coding transcripts.

We asked whether transcript properties that affect mRNA stability in other contexts could explain the range of destabilization in our data^32, 33^. For transcript length, codon optimality, and untranslated region (UTR) length, we calculated the correlation coefficient between the observed log_2_(fold-change) of the transcript k_deg_ and the parameter in question. Transcript length and optimal codon usage have been reported to have effects on association with RNP granules and mRNA stability respectively, but neither has correlation with stability in our data (Figure S3C&D)^34, 35^. The UTRs of mRNAs contain regulatory features, leading us to speculate that longer UTRs may correlate with differential stability – yet neither 3’ nor 5’ UTR length has any relationship with hypoxic destabilization (Figure S3E&F)^36, 37^.

Functional characterization shows that transcripts related to translation and the ribosome are exempt from destabilization under hypoxic conditions. We performed gene set enrichment analysis (GSEA) with transcripts ranked by the log_2_(fold-change) of k_deg_. GSEA identifies pathways associated with gene level measurements by calculating enrichment or depletion in the rank order of measured genes^38^. We found that 49 gene sets were enriched (FDR < 0.25) towards the unstable transcripts (Figure S3G). These gene sets span a range of functional categories including cytokine signaling, nutrient transport, and metabolism. In contrast, only 6 gene sets enrich (FDR < 0.25) toward those transcripts whose stability does not change (Figure S3H). These gene sets are all related to the ribosome or the regulation of translation. Thus, it appears that though most of the transcriptome is destabilized during hypoxia, transcripts related to protein synthesis are partially exempt from increased degradation.

Many ribosomal protein mRNAs along with other genes related to new protein synthesis possess a 5’ terminal oligopyrimidine (TOP) regulatory motif that connects those transcripts to mTORC1 control by way of the protein LARP1^39^. Filtering our data against a consensus list of TOP mRNAs, we find that TOP containing transcripts have a mean increase in decay rate that is 68% of all others measured (Figure 3H). We conclude that transcripts that are coregulated to control protein synthesis are partially protected from global degradation of the transcriptome, possibly through the 5‘ TOP which has been shown to stabilize transcripts elsewhere^40^.

Macrophages in hypoxia appear to compensate for increased mRNA degradation with increased transcriptional output (Figure 3E). Genes induced by LPS and IFNγ, including IL1B and CXCL9, exhibit increased synthesis to the extent that they are expressed at even higher levels than in normoxic inflammation. Utilizing the differential synthesis data in the same enrichment analysis as above, we find that SPI1, CREB1, and RUNX1 targets are enriched (Figure 3F). SPI1 has been associated with myeloid cell cytokine expression and inflammatory signaling in hypoxia, while CREB1 and RUNX1 are both reported to cooperatively bind DNA with HIF1a^41–44^.

Our results show that hypoxia does not block the macrophage inflammatory response but rather leads to widespread transcript turnover and compensatory transcriptional increase relative to activation in normoxia. However, there is significantly altered expression for about 1500 genes demonstrating hypoxia is changing gene expression during macrophage activation.

### Hypoxia alters the specificity of macrophage phagocytosis

To determine if the different gene expression in hypoxic macrophages affected specific biological functions, we performed GSEA on differentially expressed genes in hypoxia. We found that two categories were significantly depleted: gene sets related to phagocytosis and gene sets related to the ECM (Figure 4A). The most upregulated pathways are uniformly related to monosaccharide processing, consistent with an increase in glycolytic metabolism in hypoxia (Figure S4A)^45^. Downregulated gene sets related to phagocytosis included the lysosome and iron metabolism (Figure 4A, purple highlights). By separating these genes from our data and plotting them against the differential degradation and synthesis of their transcripts we see that their downregulation is largely driven by increased transcript decay (Figure 4B). Among the most affected genes in this class are TFRC, the transferrin receptor, LGMN, which has a role in antigen processing, and MSR1 and CD68 which are idiosyncratic macrophage phagocytosis receptors^46–48^. In contrast, CD14 expression moderately increases in hypoxia while TLR4 expression remains unchanged (Figure S4B).

**Figure 4.**
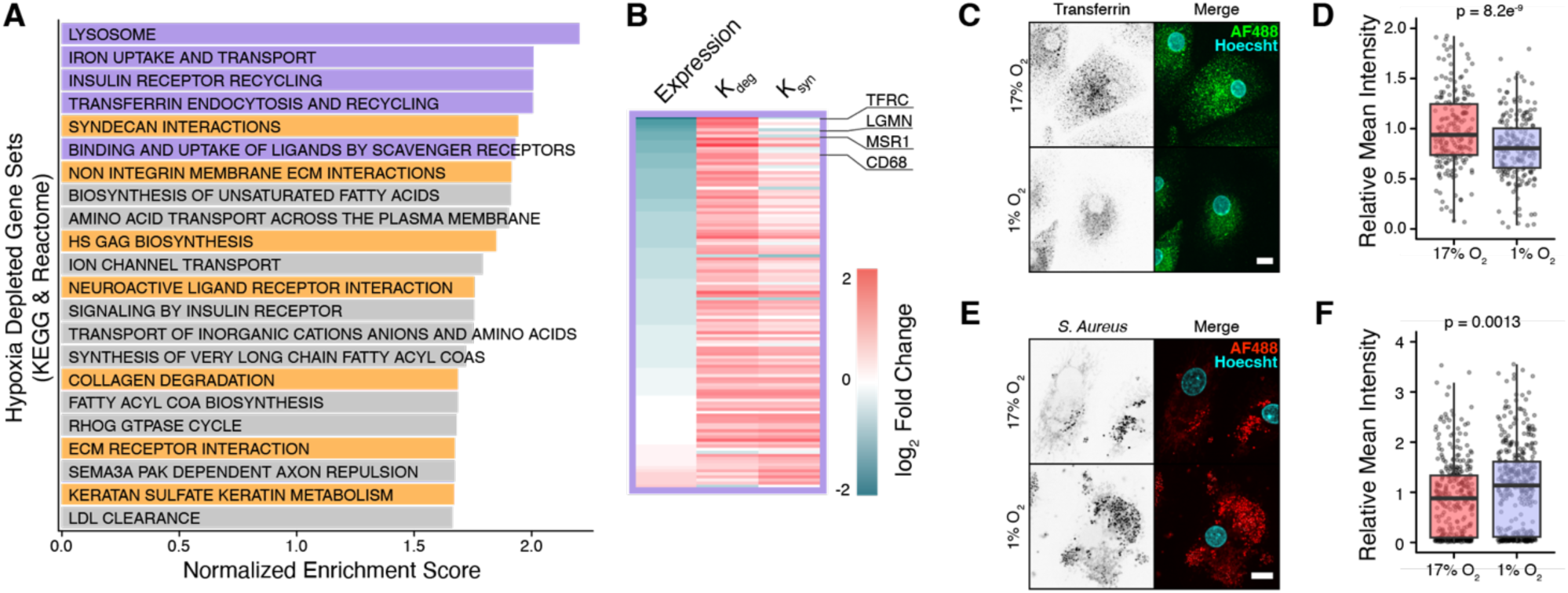
Hypoxia modifies the specificity of macrophage phagocytosis towards decreased nutrient uptake and increased pathogen ingestion. A) Results from GSEA on downregulated genes in hypoxia. Data were compared to the Kyoto Encyclopedia of Genes and Genomes (KEGG) and Reactome canonical pathways gene sets. All gene sets with FDR < 0.25% are shown. Gene sets related to phagocytosis are highlighted in purple, those related to the extracellular matrix are highlighted in orange. B) Phagocytosis related genes plotted for differential expression, differential degradation (k_deg_) and differential synthesis (k_syn_). Genes are plotted in order of ascending log_2_(fold-change) of total expression. C&E) Phagocytosis of Alexa-488 labeled transferrin (C) or pHrodo red labeled *S. Aureus* bioparticles (E). Scale bar = 10 μm. D&F) Quantification of C &E. Points represent mean fluorescence intensity of the phagocytosis channel in each cell, normalized to the mean of control intensity for each replicate (n = 3). Box plot shows median with the interquartile range and whiskers at 95% of that range. P value calculated from Mann Whitney rank sum test.

Differences in cell surface receptors associated with uptake of different ligands prompted us to hypothesize that the specificity of phagocytosis might be altered by hypoxia. Specifically, increased CD14 expression would promote the uptake of Gram-negative bacteria while the intake of transferrin would be reduced with less transferrin receptor.

We tested the specificity of uptake by hypoxic activated macrophages by supplementing their media with fluorescently tagged compounds for 90 minutes at the end of the 24-hour activation period: Alexa488-transferrin and pHrodo Red labeled, inactivated *Staphylococcus aureus* particles. Consistent with the changes in mRNA levels, we observed a 13% decrease (p = 8.2e^-9^, Mann-Whitney) in transferrin uptake by macrophages after 24 hours of induction with LPS and IFNγ in hypoxia compared to the same induction in normoxia (Figures 3A, 4C&D). With *S. aureus* we observed the opposite effect, where macrophages increased their uptake of particles by 25% (p = 0.0013, Mann-Whitney) in hypoxia relative to normoxic inflammation. This can be attributed to the increase in CD14, which has a key role in the recognition and phagocytosis of Gram negative bacteria^49^.

Overall, we conclude that hypoxic, activated macrophages reprioritize the specificity of phagocytosis by altering the gene expression of internalization receptors. We speculate that the tuning of specificity could be a way of prioritizing the clearance of pathogens when oxygen availability is low.

### Hypoxia reduces macrophage digestion of the ECM and tissue invasion

Gene sets related to ECM interaction and digestion are also depleted in hypoxia (Figure 3A, orange highlight) due to increased mRNA degradation (Figure 5A). Among the most affected genes are C5AR1, a receptor which regulates hyaluronic acid production, metalloproteinases (i.e. MMP7 and MMP9), integrins (i.e. ITGA6, ITGB8), and SDC2 encoding syndecan-2, a proteoglycan that mediates interaction with the ECM^50–52^.

**Figure 5.**
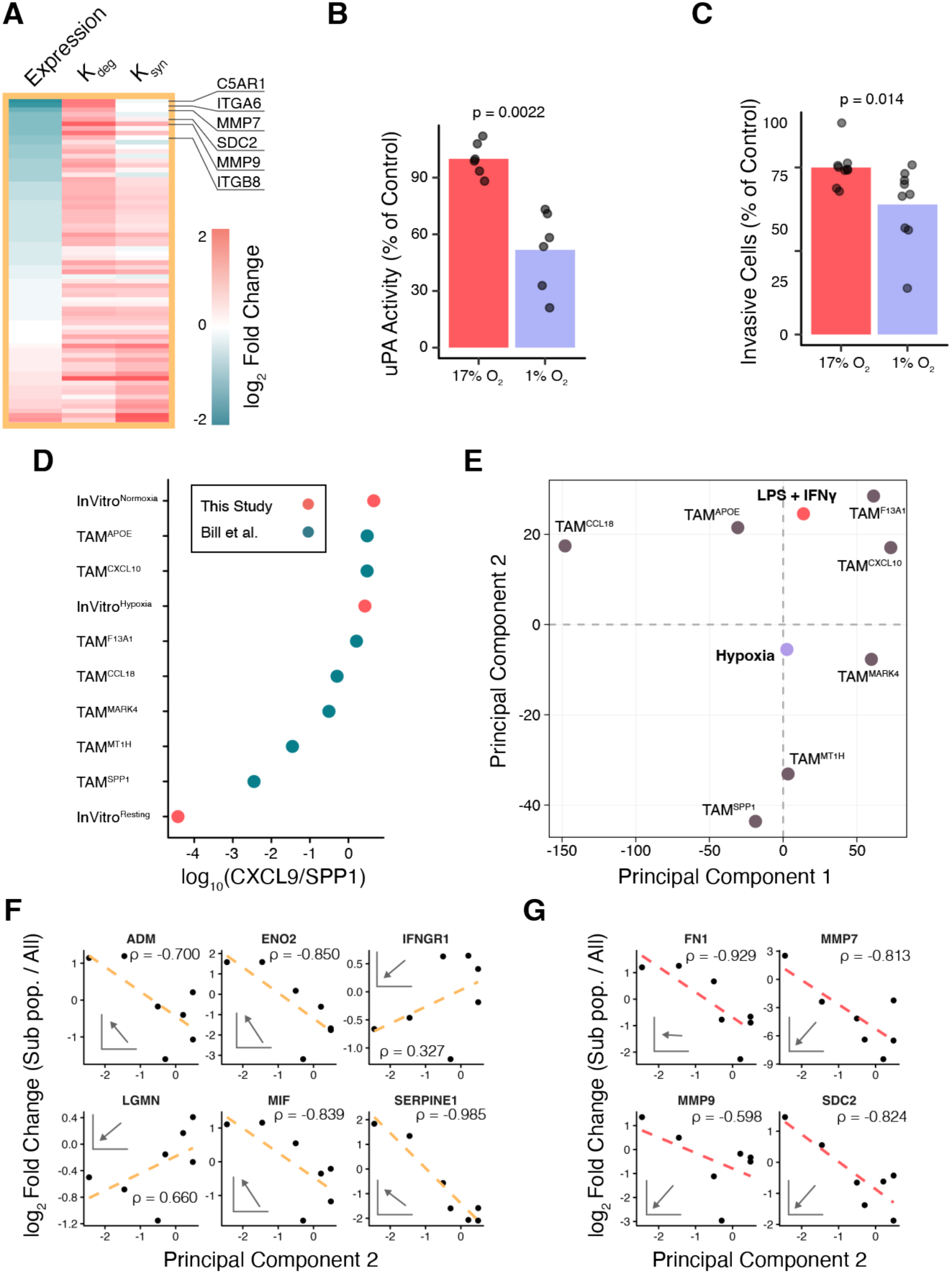
Hypoxia reduces tissue invasion and confers a tumorigenic phenotype in macrophages. A) ECM related genes plotted for differential expression, differential degradation (k_deg_) and differential synthesis (k_syn_). Genes are plotted in order of ascending log_2_(fold-change) of total expression. B) Macrophage supernatant uPA activity assay where measurements are normalized to the mean of control. P value calculated from Mann Whitney rank sum test, n =6 for each condition. C) Transwell assay where the percentage of cells passing through an ECM coated filter was normalized to the control mean of each biological replicate. P value calculated from Mann Whitney rank sum test, where total n = 9 with three biological replicates and three technical replicates each. D) log_10_ of the CS ratio of tumor associated macrophage (TAM) subtypes from Bill et al. and bulk RNA seq from this study E) Principal component analysis of relative gene expression among macrophage subpopulations from Bill et al with the log_2_(fold-change) from this study projected into those coordinates. F) Example genes that correlate with the second principal component of subpopulation relative expression. Correlation coefficient ρ is the calculated Pearson coefficient for those genes. The vector drawn on the graph is the unit vector of that gene in log_2_(fold-change) space of the hypoxia to normoxia comparison of macrophage gene expression in this study. G) Example genes with correlation to the second principal component as above but with low or no agreement with the unit vector of gene expression from this study.

Modulating the expression of ECM components and interactors suggests that in hypoxia, macrophages are less prone to tissue invasion and less active in matrix remodeling. Earlier we had noted that one of the transcripts most impacted by hypoxic mRNA instability is PLAU (Figure 3C&D). Regulation of plasminogen is a key component in tissue remodeling by macrophages, and thus we decided to confirm whether hypoxia does indeed alter plasminogen activation activity of hypoxic activated macrophages^30, 53^.

We hypothesized that the activity of uPA, the protein product of PLAU, would decrease in hypoxia because its mRNA expression is reduced and expression of a direct inhibitor, serpin E1, is increased in the supernatant from cultured hypoxic macrophages (Figure S5A). In an assay of supernatant uPA activity, we found that hypoxia results in a 43% reduction (p = 0.0022, Mann-Whitney) in activity after 24 hours (Figure 5B). We interpret this result to be the integration of decreased PLAU mRNA expression and increased serpin E1 inhibition.

The plasminogen system is one of several ECM protease pathways with reduced expression in hypoxia, which prompted us to examine whether hypoxic activated macrophages have an overall impairment of ECM invasion compared to normoxic activation. We assessed whether hypoxia affects the activated macrophage capacity to invade a layer of ECM in a transwell assay. We found that over 24 hours of hypoxia 15% fewer (p = 0.014) cells were able to traverse the membrane in our assay. We conclude that overall gene expression changes in hypoxic macrophages lead to a decrease in invasive migration by way of ECM remodeling.

Together we conclude that hypoxia reduces inflammatory macrophage capacity to invade tissue matrix due to downregulation of proteases and ECM receptors that would enable them to digest, reorganize, and traverse through the ECM. Such a shift suggests that activated macrophages reduce their motility when they encounter hypoxic environments.

### Hypoxia drives macrophages towards a tumorigenic phenotype

The signaling status of macrophages impacts the surrounding tissue, and dictates how they respond to the environment, and so we examined how hypoxia alters the signaling profile of activated macrophages. We examined the differential expression of cytokines and cytokine receptors between normoxic and hypoxic activated macrophages (Figure S5B&C). We found CXCL11, IL1A, IL1B, and other canonical inflammatory signals are upregulated^54^. The PPAR receptors have differential expression based on the phase of inflammation^55^. In hypoxic activation PPARG is upregulated while PPARA is slightly downregulated compared to normoxic activation, suggesting that hypoxic activated macrophages have shifted towards a resolution of inflammation phase by our 24-hour timepoint. Angiogenic factors ADM, VEGFA, VEGFB, and receptors FLT1 and ACVRL1 are also upregulated in hypoxic activation. Angiogenesis pathways are known to be activated by hypoxia, but we found it notable that they could coexist with upregulation of inflammatory pathways^56^.

Changes in hypoxic macrophage signaling prompted us to ask whether any of these changes inform on how hypoxia affects macrophages that have entered a solid tumor. A recent study of gene signatures in 52 different human tumor biopsies found that the ratio of CXCL9 and SPP1 (CS) serves as an effective biomarker for tumor response to treatment and correlates with a range of pro-tumorigenic (CS^high^) or anti-tumorigenic (CS^low^) pathways^7^. We compared the macrophages in our study to these tumor associated macrophages to assess how hypoxia influences gene expression in the context of a solid tumor.

In our data, resting macrophages (CS = 3.8 e^-5^) and normoxic inflamed macrophages (CS = 4.4) define the endpoints of the CS range (Figure 5D). This places the resting and inflammatory states at opposite ends of the CS spectrum, with hypoxic inflammatory macrophages shifted slightly toward the middle (CS = 2.6) of the subpopulations analyzed by Bill et al^7^. Macrophage polarity defined as a ratio of these two biomarkers is a new concept, so we reanalyzed their dataset to gain additional insights into how hypoxia relates to the diversity of tumor associated macrophages.

We began by performing a principal component (PC) analysis on the subpopulations of cells reported in Bill et al. We chose PC analysis because it would allow us to transform expression data of tumor macrophage sub populations into a vector space where we can distinguish biologically meaningful clusters from one another and extract the most influential genes. Our goal was to define a tumor associated macrophage PC space independent of our data, into which we could project our results and draw conclusions about the effects of hypoxia and activation.

Starting with the data from Bill et al. filtered down to only macrophage subpopulations analyzed in their study, we further filtered the count matrix by applying a threshold of 1000 read counts per gene in the sum of all macrophages. The relative differential expression of transcripts for each sub population was calculated by dividing the unitized read count vector of each sub population by the unitized, summed read count vector for all macrophages. These relative expression values in log_2_(fold-change) units were then used for PC analysis (Figure S5D).

The analysis shows that while the majority of the variance is captured by the first dimension of PC space, that dimension has low specificity: it is comprised of minor contributions of many genes. The second dimension is highly influenced by only a few genes (Figure S5E&F). Further, this second dimension best recapitulates the CS ratio among the subpopulations (Figure S5D, point color scale).

We projected the log_2_(fold-change) of our data into the PC space defined by the single cell RNA-seq tumor macrophages. The gene expression changes associated with activation from resting to inflammatory macrophages in normoxia clustered with the sub populations with the highest CS ratio. The gene expression changes between activated normoxia and activated hypoxia trend towards the low CS ratio subpopulations and were most similar to the MARK4-enriched sub population (Figure 5E). This suggests that inflammatory activation by LPS and IFNγ generates a gene expression profile most like the CS^high^ tumoricidal macrophages. Conversely, hypoxia in activated macrophages drives a gene expression profile in the direction of CS^low^ tumorigenic macrophages.

Seeking to gain a better understanding of the gene specific behavior of the PC space, we asked which pathways were associated with the second dimension because it best recapitulated the CS ratio. We performed GSEA on the relative contributions of genes to the axis (Figure S5G). Pathways associated with negative values of the second PC dimension, (CS^low^, tumorigenic), include translation, translation regulation, nonsense mediated decay of mRNA, and the response to starvation. Pathways associated with the positive values of the second PC dimension (CS^high^, tumoricidal), include antigen processing, interferon gamma signaling, and the lysosome. The enrichment of these pathways shows that the second PC dimension is defined by functional groups of genes with striking similarity to those identified by our dataset comparing activated normoxic macrophages and activated hypoxic macrophages.

We were curious whether specific genes from our TimeLapse-seq results correlated with the second dimension of PC space and were thus drivers of the pathways identified by GSEA. When we compared our dataset to the second dimension of PC space on a gene-by-gene basis we found that hypoxia marker gene ENO2 has a negative correlation with the inflammatory axis and agrees with our data (Figure 5F). This was also the case for MIF and ADM, genes associated with angiogenesis, and for SEPRINE1, the gene encoding the inhibitor of uPA protease activity. IFNGR1, part of the IFNγ receptor, and LGMN, a component of lysosomal processing, have a negative correlation with the axis, in keeping with their association with inflammation.

Not every gene correlating with the second PC dimension agrees with our hypoxic data. FN1, encoding fibronectin, the genes for metalloproteinases MMP7 and MMP9, and SDC2, encoding syndecan-2 all had strong negative correlations with the second PC dimension. However, in our experiment, hypoxia downregulates or has no association with these genes (Figure 5G).

We conclude that hypoxia drives gene expression of tumor associated macrophages toward tumorigenic subtypes in agreement with Bill et al. Our results show that hypoxia is just one component of the tumor microenvironment, and it appears that while reduced engagement of the ECM is a phenotype brought on by hypoxia and inflammatory activation *in vitro*, it is not part of the tumorigenic phenotype in these populations. Rather, hypoxia appears to reduce lysosomal activity, while upregulating genes related to translation, angiogenesis, and pathways related to RNA decay (Figure 6).

**Figure 6.**
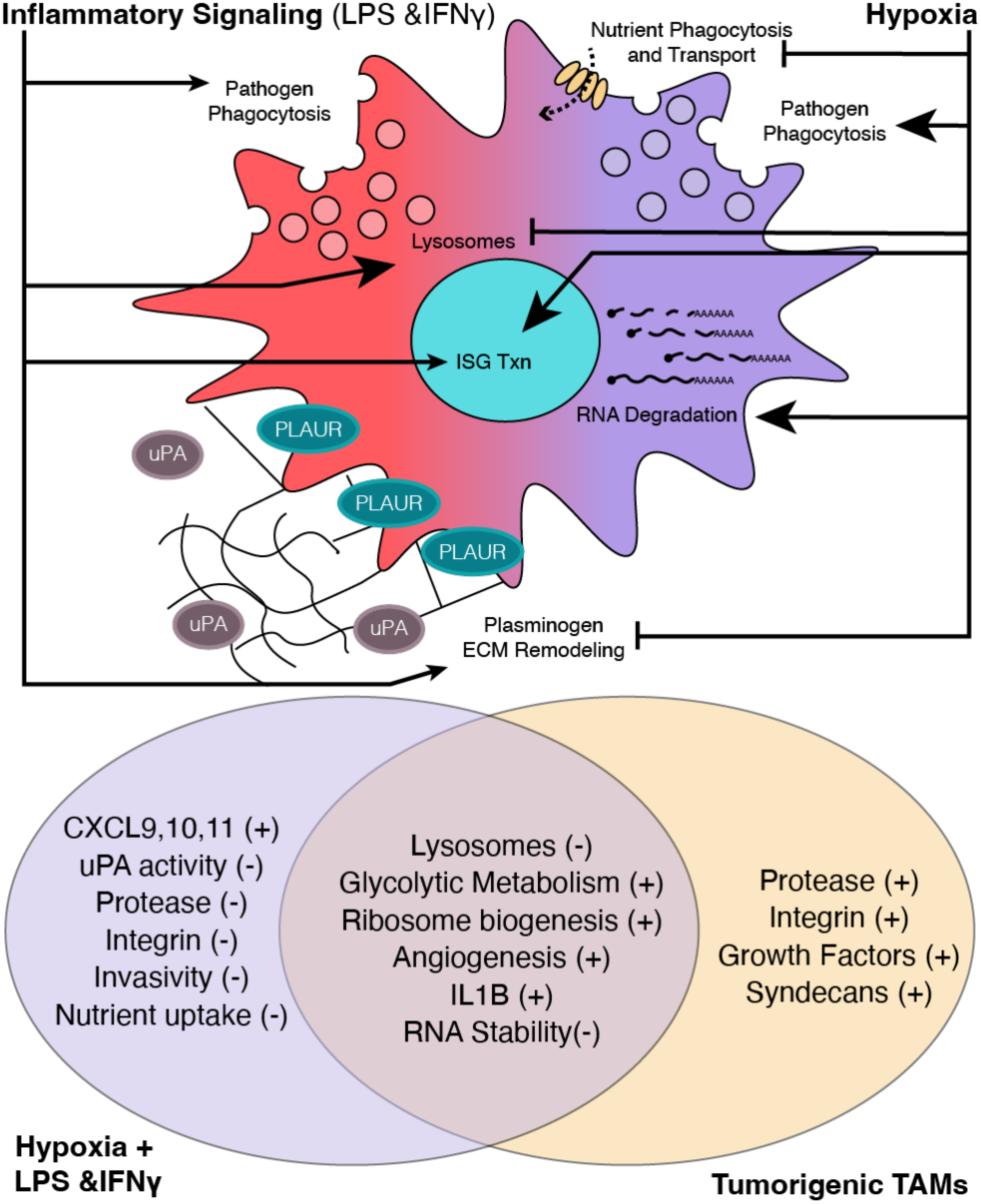
Hypoxia modifies the function output of inflammatory macrophages with similarity to tumor associated macrophages. Schematic of processes modulated by hypoxia in activated macrophages and Venn diagram of overlap with tumorigenic tumor associated macrophages. Abbreviations: ISG – Interferon Stimulated Gene, TAM – Tumor Associated Macrophage.

## Discussion

In this study we used TimeLapse-seq to comprehensively measure the dynamics of the transcriptome during macrophage activation in normoxia and hypoxia. We showed that hypoxia induces a global shift toward mRNA transcript instability, though transcripts encoding ribosomal proteins with 5’ TOP motifs appear to be exempt to some degree. The functional ramifications of hypoxic transcriptome reprograming *in vitro* are the altered specificity of phagocytic uptake by macrophages and the reduction of proteolytic digestion of the ECM. Tumor associated macrophages that are more tumorigenic are associated with hypoxia and share the reduced expression of lysosomal factors, increased angiogenic output, and increased expression of ribosomal proteins.

These findings are most directly related to two scenarios of tissue regulation/dysregulation: wound healing and solid tumors.

Critical to proper wound healing, macrophages are well adapted to maintain their inflammatory response programs in the face of decreased oxygen levels. Certain inflammatory cytokines (i.e. IL1A&B, CXCL11) are amplified in hypoxia. At the same time, we also see some hallmarks of macrophages entering inflammation resolution in hypoxia (i.e. PPARG), reflecting the role of hypoxia as a signal to initiate wound healing^55^. Our dataset provides insight into gene expression regulation in effector myeloid cells like macrophages during the transition from an initial inflammatory phase into a regenerative one. To do so, macrophages must regulate clearance of damaged tissue, removal of pathogens, and initiation of regenerative processes such as angiogenesis simultaneously^57, 58^. We speculate that increased mRNA turnover coupled with increased rates of transcription facilitate the rapid activation and deactivation of these pathways in response to stimuli from the environment.

The altered phagocytic function of hypoxic macrophages raises specific issues related to the clearance of pathogens in a wound healing context. Many pathogens are phagocytosed by macrophages but then continue to proliferate intracellularly, thereby evading further immune system response^59^. Examples include *Leishmania*, *Yersinia*, and *Mycobacteria*^60–62^. Lesions formed by these pathogens are often hypoxic, and we hypothesize that the downregulation of lysosomal machinery observed as a consequence of hypoxia may help to explain why macrophages phagocytose these pathogens but then fail to kill them^63, 64^.

Likewise, hypoxia in tumors appears to induce macrophages to express pathways that would be beneficial in other contexts but are coopted by the tumor to promote growth. These are not new ideas – tumors have long been associated with hypoxia, and tumor associated macrophages have been shown to express angiogenic signals that promote tumor growth^8, 65^. Our study adds a new dimension to this dogma: a macrophage that receives both inflammatory signals like IFNγ and hypoxia will still be induced to produce more regenerative signals. Furthermore, tumor associated macrophages would be subjected to increased mRNA degradation, perhaps further degrading the inflammatory gene program over time. Our study captures dynamics only at 24 hours, but future work may reveal how hypoxic inflammatory macrophages develop over time to become fully tumorigenic.

The reduction in macrophage ECM interaction and invasion during hypoxia is notable because previous work has associated increased expression of motility factors and ECM proteases with tumorigenesis^66, 67^. Indeed, the tumorigenic macrophage populations from Bill et al. were expressing these pathways at high levels. Our work suggests that either hypoxia alone does not induce these pathways in tumors, or that inflammatory activation is enough to drive gene expression of these factors in the opposite direction.

Two other studies have observed widespread hypoxic mRNA decay in generic tissue culture models, supporting our findings in primary human macrophages^68, 69^. Using conventional mRNA lifetime calculations from actinomycinD transcription assays and a s^4^U pull-down approach in HUVECs, Tiana et al. find that decay rates are about 1.6 times faster in hypoxia for a dataset of a few hundred genes^69^. In a study of Thp1 cells, an immortalized leukemic monocyte line, Bauer et al. find nearly 600 mRNAs to be differentially degraded using SLAM-seq, a metabolic labeling approach similar to TimeLapse-seq^68, 70^. Our data, produced with TimeLapse-seq and analyzed with bakR, identify thousands of transcripts regulated by post-transcriptional decay in primary human macrophages. Together these three studies point to global transcriptome decay in hypoxia as a fundamental regulatory event.

Hypoxic mRNA instability appears be governed by mTOR regulation. Tiana et al. describe a mechanism by which TSC2, an mTOR regulator, is necessary to induce hypoxic mRNA instability. Our findings suggest that 5‘ TOP motif mRNAs are partially exempt from hypoxic decay, but what mechanisms act to exempt TOP RNAs from global degradation remain unknown. Some the transcripts which are most highly degraded in hypoxia (i.e. TFRC, PLAU) have well studied 3’ UTR regulatory motifs that are known to modulate transcript levels in various conditions^71, 72^. We anticipate that RNA binding proteins will play a major role in determining which transcripts are either targeted for or protected from hypoxic RNA degradation, though whether these rely on general principles in all cells or are idiosyncratic to specific lineages remains to be seen.

We conclude that hypoxia is an important environmental cue of macrophage phagocytic and migratory behavior and that fundamental post-transcriptional regulation mechanisms drive those changes. Further study to reveal the specific post-transcriptional mRNA pathways at work will yield a deeper understanding of how global shifts in mRNA expression and stability contribute to the specific cellular function of macrophages and other innate immune cells.

## Acknowledgements

We would like to thank Matthew Kwan and all members of the Parker Lab for their assistance with preparing the manuscript. We thank Isaac Vock and Meaghan M.S. Courvan for advice and guidance on analysis. The work was funded by the Howard Hughes Medical Institute and the Damon Runyon Cancer Research Foundation.

## Author Contributions

E.M.C.C. performed all experiments and analysis. Both authors conceptualized the study and prepared the manuscript.

## Data and Code Availability Statement

All original sequencing datasets will be made available in GEO on final publication. Analysis scripts will be made available at https://github.com/ParkerLabCU.

## Methods – Experimental Procedures

### Macrophage Culture Conditions

Unpolarized primary human peripheral blood macrophages were purchased from StemCell Technologies (Cat #70042). Sequencing experiments were conducted with donor and lot# matched cells. Macrophages were cultured in ImmunoCult SF Macrophage Medium (StemCell Technologies Cat. #10961). For inflammatory activation, macrophages were induced with media supplemented with 5 μg/mL eBioscience lipopolysaccharide (Thermo Fisher, Cat. # 00-4976-93) and 50ng/mL interferon gamma (StemCell Technologies, Cat. #78141.1). Normoxic samples were cultured in a 37°C humidified standard tissue culture incubator in 5% CO_2_ mixed with room air which is equivalent to about 17% O_2_ at sea level with an O_2_ partial pressure of 120 mmHg. Hypoxic samples were cultured in an atmosphere-controlled Oxford Optronix HypoxyLab incubator set to 7 mmHg O_2_ (1% at sea level), 5% CO_2_, 37°C, 70% relative humidity, and balanced with nitrogen. Media for hypoxic samples was preconditioned for two hours before cells were transferred into the incubator and media changed.

### TimeLapse-seq

TimeLapse-seq^16^ requires a s^4^U feed to metabolically label newly transcribed RNA. Metabolic labeling was performed by supplementing s^4^U (Sigma, Cat. # T4509) dissolved in water to a final concentration of 500 μM for the last two hours. Macrophages were scraped from culture dishes and resuspended in ice cold PBS. Cells were then immediately pelleted and lysed into Trizol (Thermo Fisher, Cat. # 15596026) in LoBind Tubes (Eppendorf, Cat. # 022431021). MTS-TAMRA (Biotium, Cat. # 91030) control labeling was performed using the reaction conditions outlined in Duffy & Simon 2016^73^. For preparation of TimeLapse libraries, RNA was reduced by adding 1 mM dithiothreitol (Thermo Fisher, Cat. # D1532) following Trizol extraction. Genomic DNA was removed from samples using Turbo DNase (Thermo Fisher, Cat. # AM2238). Oxidation chemistry was performed by adding RNA to a solution of 100 mM sodium acetate (Thermo Fisher, Cat. # AM9740), 4 mM EDTA (Thermo Fisher, Cat. # AM9260G), 5% v/v 2,2,2-Trifluoroethylamine (Sigma, Cat. # 269042), and 10 mM sodium periodate (Thermo Fisher, Cat. # 419610050) and incubating at 45°C for 1 hour. RNA was then bead purified using RNAClean beads (Beckman Coulter, Cat. # A63987) and reduced in 1 mM dithiothreitol, 10 mM Tris pH 7.0 (Thermo Fisher, Cat. # AM9850), 1 mM EDTA, and 100 mM NaCl (Thermo Fisher, Cat. # AM9760G) for 30 min at 37°C. RNA was purified and concentrated with RNAClean beads before proceeding with library preparation.

### Library preparation and sequencing

Library preparation was carried out using the Takara SMARTer Stranded Total RNA-Seq Kit v3 – Pico Input, Mammalian (Cat. # 634485). Prep was performed exactly as the protocol states with the following parameters. RNA was chemically fragmented for 90 seconds before proceeding with First-Strand cDNA synthesis. PCR step 1 was performed with 10 cycles of amplification, PCR step 2 was performed with 9 cycles. All samples were assessed for library quality using an Agilent Bioanalyzer. Sequencing was performed on a single lane of an Illumina NovaSeq Paired End 150 platform sequencer by Novogene. Raw read depth was an average of 3.8 million reads per experimental sample.

### Supernatant cytokine content and uPA activity

Macrophage supernatants were evaluated for their cytokine content using a Human Cytokine Array Kit (R&D Systems, Cat. # ARY005B). Supernatants were evaluated in biological duplicate. Enrichment or depletion is reported for assay spots that reproduce with greater than 2-fold-change over normoxic control. uPA content was measured using a Urokinase-type Plasminogen Activator Chromogeneic Activity Assay Kit (Abcam, Cat. # ab108915). Chromatic read out was performed on a BMG Labtech CLARIOstar Plus microplate reader.

### Transwell invasion assay

Macrophage invasion was evaluated using a Boyden chamber transwell assay. Chambers were set up in 12 mm diameter Millicell Cell Culture Inserts with a polycarbonate membrane with 8.0 μm pore size (Millipore Sigma, Cat. # PI8P01250) where the pores were filled with a layer of ECM from Engelbreth-Holm-Swarm murine sarcoma (Sigma, Cat. # E1270). Macrophages were applied to the upper chamber in media supplemented with IFNγ and LPS and allowed to migrate through the membrane over 24 hours in either normoxic or hypoxic atmosphere. After incubation, wells were fixed with 4% Paraformaldehyde in PBS (Santa Cruz, Cat. # sc-281692) and stained with 1 μg/mL Hoechst 33258 (Thermo Fisher, Cat. # H3569). Whole wells were imaged in wide field using a 10X objective and image stitching. Cells on the upper layer were then removed and each well was imaged again.

### Phagocytosis Assays

Macrophages were plated in 96-well plates with an optical polycarbonate coverslip base (Ibidi, Cat. # 89626) in the conditions described above. In the last 90 min of incubation, media was supplemented with either pHrodo red *S. aureus* BioParticles (Thermo Fisher, Cat. # A10010) or Human Transferrin conjugated to Alexa Fluor 488 (Thermo Fisher, Cat. # T13342). Supernatant was removed, cells were washed with PBS and then fixed with 4% PFA in PBS and stained with 1 μg/mL Hoechst. Cells were then imaged using a 60X objective and spinning disk confocal mode.

### Imaging Configuration

Imaging in this study was carried out with a Nikon dual wide-field/CSU-W1 SoRa Spinning Disk Confocal setup. The objectives used in this study were CFI60 Plan Apochromat Lambda 10X Objective, N.A. 0.45, and a CFI SR Plan Apo IR 60X Water Immersion Objective, N.A. 1.27. Wide field illumination uses a SOLA Light Engine GEN III. Confocal illumination uses a Nikon LUNF-XL 6-line laser launch equipped with 405, 445, 488, 514, 561, and 640 nm lasers. The detection setup used was a Hamamatsu ORCA Fusion BT sCMOS camera.

## Methods – Data Analysis

### Analysis of raw TimeLapse data

Analysis of sequencing reads was performed using the same pipeline described in Vock & Simon, 2023^20^. Briefly, reads were filtered for unique sequences using FastUniq v.1.1 and trimmed of their adaptor sequences with Cutadapt v.2.3. Alignment was performed using HISAT-3N using the current version of GRCh38 from NCBI (GCA_000001405.29). Uniquely mapping reads were quantified using HTSeq v.0.9.1. Aligned reads were prepared for mutational analysis with the bam2bakR workflow.

### Differential expression, stability, and synthesis analysis

The output of the preprocessing pipeline was used as count tables for DESeq2 and bakR to generate differential expression and differential stability results. Raw count tables were filtered for genes with more than 100 counts total across all samples. Differentially expressed genes called as those with P_adj_ > 0.05. Differential stability was determined by running bakR with the hybrid model using default parameters and calling those genes where P_adj_ > 0.05. These results were then used to determine differential synthesis as described in the documentation for bakR. The log_2_ fold-change of the synthesis rate was simply derived as the log_2_ fold-change of expression plus the log_2_ fold-change of the stability. Standard error of that parameter was calculated as the square root of the sum of squares of the standard errors of fold-change expression and stability. A P-value is obtained using the asymptotic Wald test and then adjusted with the Benjamini-Hochberg procedure.

### Enrichment Analysis

Enrichment analysis was performed using two tools – Enrichr and the classic Gene Set Enrichment Analysis (GSEA) software v.4.3.1^38, 74–76^. Enrichment of transcriptionally upregulated genes was performed in Enrichr against the ENCODE and ChEA consensus transcription factor gene set library. Enrichment of differential expression and stability data was carried out by generating a ranked order of enrichment and depletion for each experiment analyzed. That ranked list was used as input for the pre-ranked mode of the GSEA tool with 2000 permutations where each dataset was compared against the KEGG and Reactome canonical pathway lists of gene sets in the Molecular Signatures Database.

### Transcript destabilization classification

The Ensembl GRCh38 genome annotation version 110 was used to classify transcript type, UTR length and gene length. For genes producing multiple isoforms, the dominant annotated dominant isoform was used. Coding sequence codon optimality was determined using the Codon Statistics Database^77^. TOP motif transcripts were categorized using the consensus list determined by Philippe et al^39^. Genes categorized as expressing cytokines or cytokine receptors were derived from gene lists found at ImmPort.org^78^.

### Comparison of TimeLapse-seq data to single cell RNA-seq

TimeLapse-seq datasets were compared to single cell RNA-seq data from Bill et al. using a principal component approach. First, the full tumor dataset was filtered down the macrophage subsets described in the original paper. Low expression genes were filtered out by applying a threshold of 1000 counts per gene summed over all macrophage subtypes. The filtered dataset was then used to compute a relative expression profile for each subtype. First, a sum of all macrophages was computed and then the sum and each subtype were unitized. Relative expression was defined as the log_2_(subtype expression unit vector / summed expression unit vector). Principal component analysis was performed on the six subtypes. Differential expression values from TimeLapse-seq as log_2_(fold-change) was then projected into the principal component space by multiplying the loadings of each gene into each principal component by the corresponding gene in the TimeLapse-seq dataset. GSEA was performed on genes ranked by the loading coefficient of the second principal component. The correlation between the second principal component and the relative expression was computed as a Pearson coefficient (ρ).

### Image analysis

Transwell assay analysis was carried out by first setting all pixels outside the well membrane plus a 100-pixel buffer to zero. Nuclei were segmented using the built-in “robust background” method in Cell Profiler which sets a threshold based on two standard deviations above the mean pixel value. Nuclear area was used as a proxy for the number of cells present in the field of view to avoid error due to the erroneous merge or fractioning of closely spaced or bright nuclei.

Phagocytosis was quantified by first segmenting nuclei using a minimum cross-entropy threshold and then extending those masks to the cytoplasm of cells using the blurred phagocytosed particle channel as a guide for watershed spreading. The phagocytosis channel was background corrected by subtracting the median pixel value of the image. Intensity within the cytoplasmic masks was quantified as the mean intensity in each mask for each cell.

### Software and data availability

Software versions not stated above include R v.4.3.0, Cell Profiler v.4.2.4, FIJI v. 2.3.0/1.53v. Analysis scripts are available at https://github.com/ParkerLabCU. Raw sequencing data are available from the gene expression omnibus accession #. Processed data files are available as supplementary data.

## Supplemental Figures

**Figure S1, Related to Figure 1.**
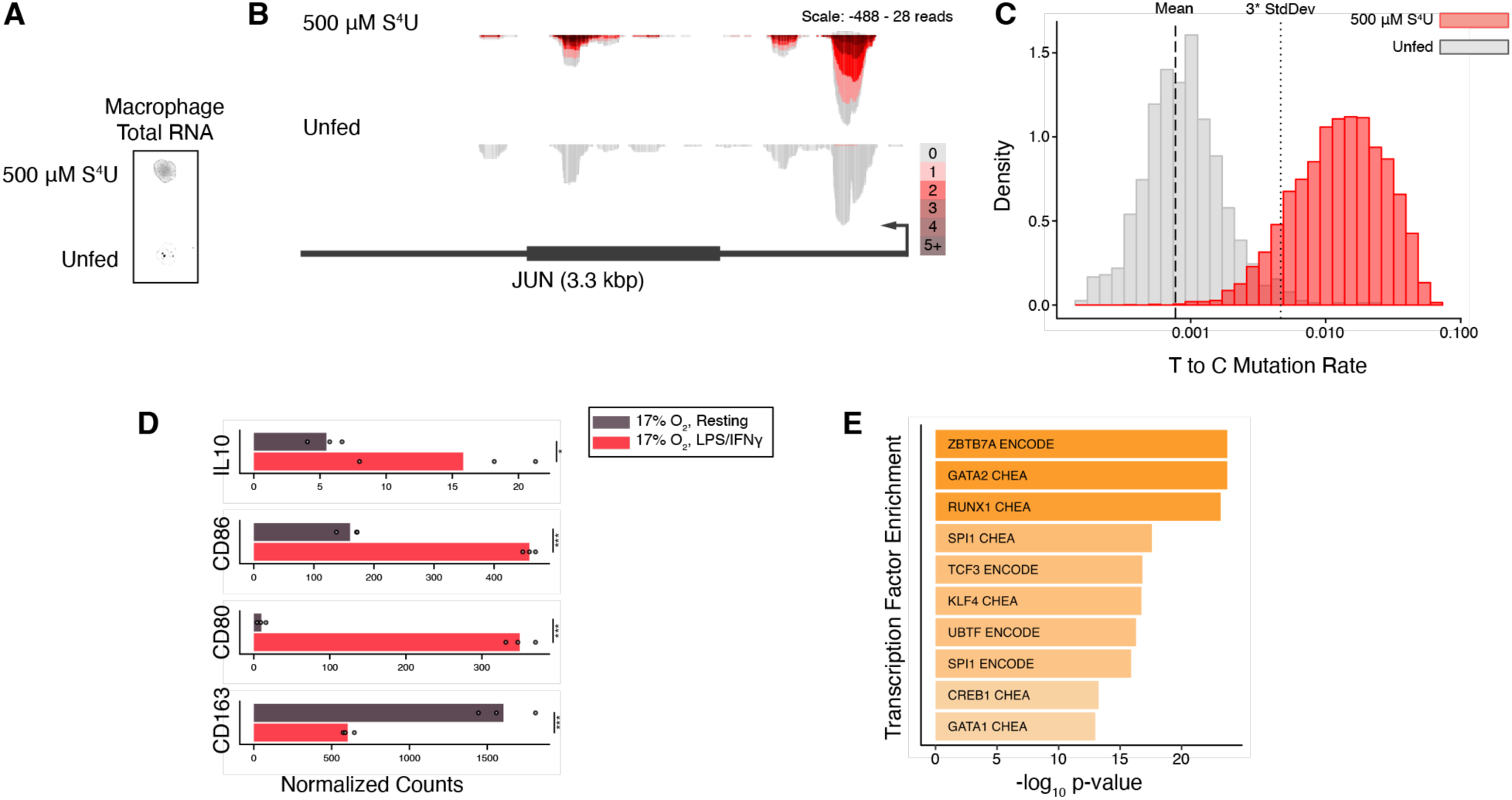
Macrophages uptake s^4^U at rates detectable by TimeLapse-seq. A) Fluorescent blot of RNA extracted from macrophages fed with 500 μM s^4^U for two hours compared with unfed control, where RNA is conjugated with an MTS-TAMRA fluorophore. B) Genome browser track for JUN showing detected mutations in s^4^U fed samples compared to unfed. Color bar corresponds to number of mutations detected per read. C) Density histogram of mutation rate per read for fed (red) and unfed (grey) samples. Annotations include the mean mutation rate of unfed samples (dashed line, 7.7 e^-4^) and three standard deviations above that mean (dotted line, 4.6 e^-3^). D) Normalized read counts for specific genes related to macrophage activation where points are individual replicates and bars represent the mean of those replicates. FDR value thresholds given as * FDR < 0.05, ** FDR < 0.01, *** < 0.001. E) Top ten transcription factors with enriched promotor binding sites for transcripts with decreased synthesis. Enrichment is drawn from the consensus database of ENCODE and ChEA transcription factors.

**Figure S2, related to Figure 2.**
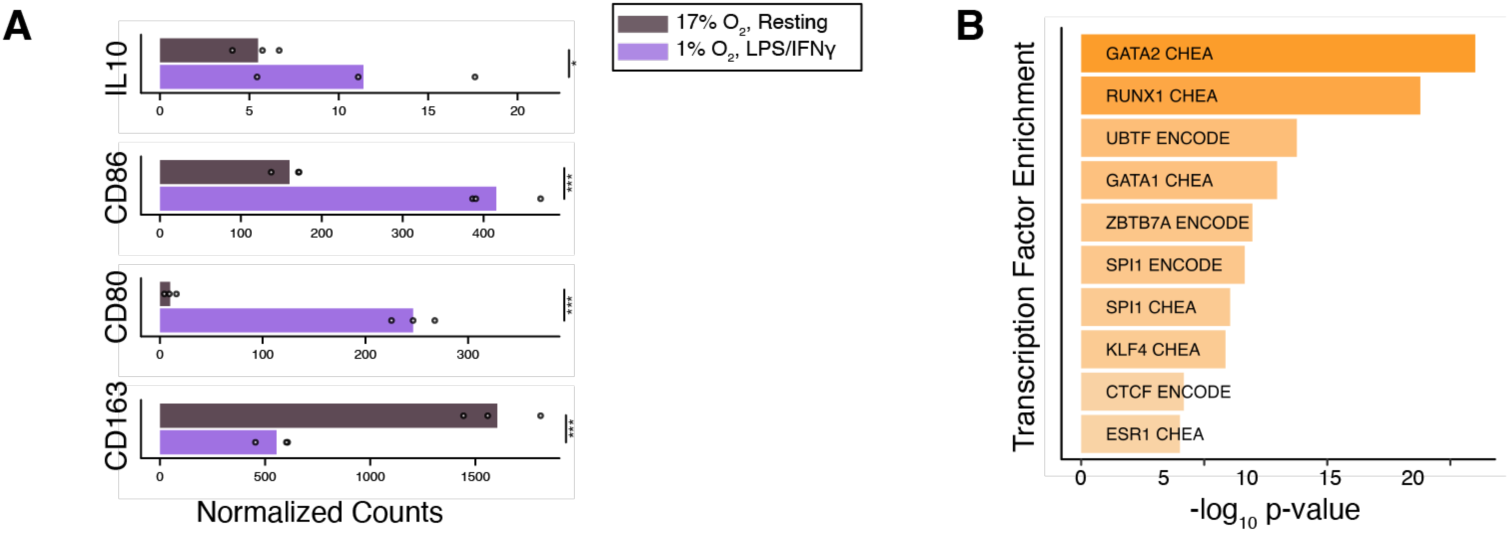
Macrophages activate in hypoxia. A) Normalized read counts for specific genes related to macrophage activation where points are individual replicates and bars represent the mean of those replicates. FDR value thresholds given as * FDR < 0.05, ** FDR < 0.01, *** < 0.001. Top ten transcription factors with enriched promotor binding sites for transcripts with decreased synthesis. Enrichment is drawn from the consensus database of ENCODE and ChEA transcription factors.

**Figure S3, related to Figure 3.**
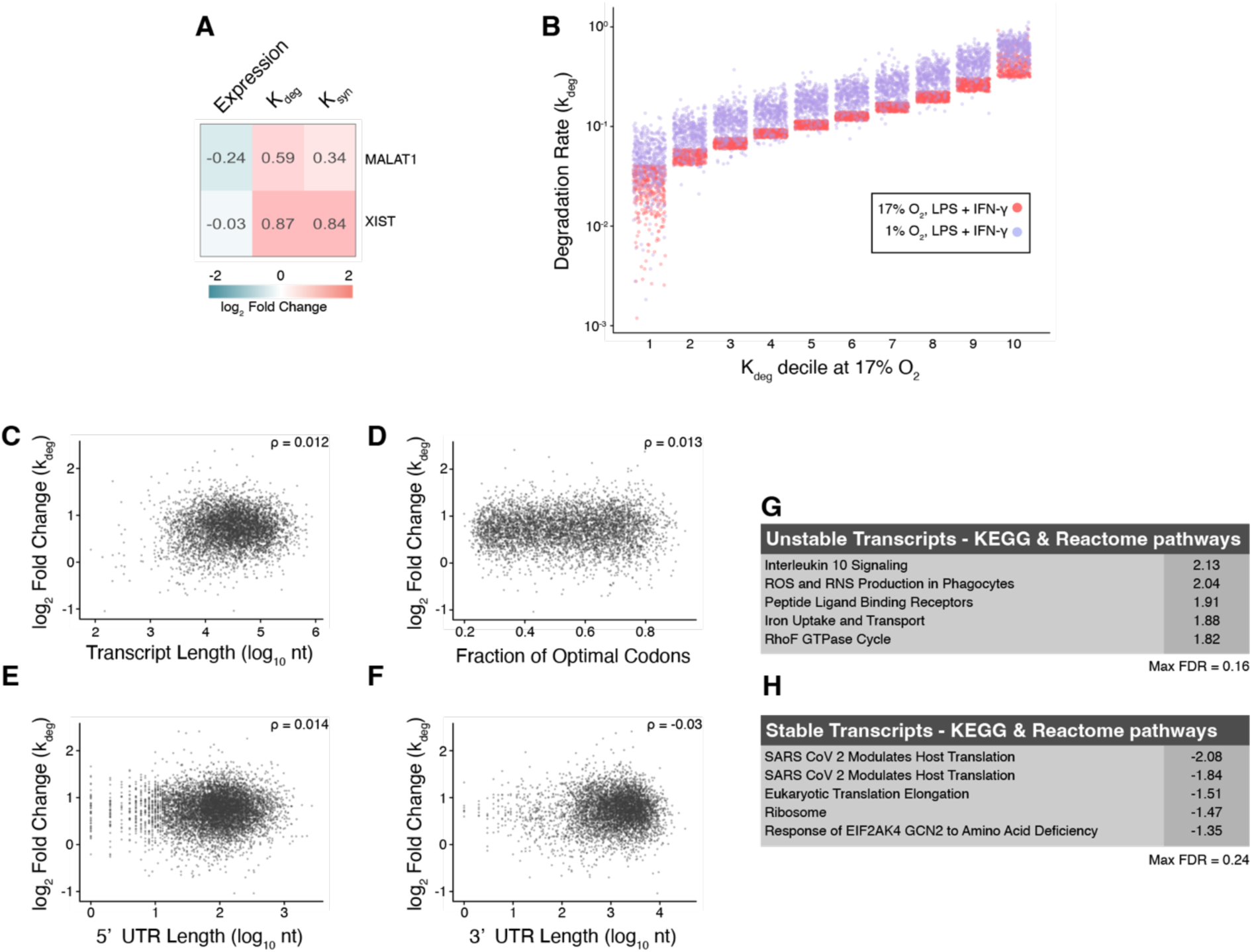
General properties of degraded transcripts do not predict differential degradation in hypoxia. A) Labeled heat map of expression levels, degradation rate (k_deg_), and synthesis rate(k_syn_) for long non-coding RNAs MALAT1 and XIST comparing activated macrophages between normoxia and hypoxia. B) Transcripts binned by decile of normoxic k_deg_ and plotted against their k_deg_ in each oxygen condition. C-F) Plot of differential degradation rate (k_deg_) dependency on C) total transcript length, D) Codon optimality, E) 5’ UTR length and F) 3’ UTR length. ρ gives the Pearson coefficient between the two parameters. G&H) GSEA of transcripts ranked by differential stability in hypoxia compared to normoxia where G) gives unstable pathways in hypoxia and H) gives stable pathways. Given values are the normalized enrichment scores.

**Figure S4, related to Figure 4.**
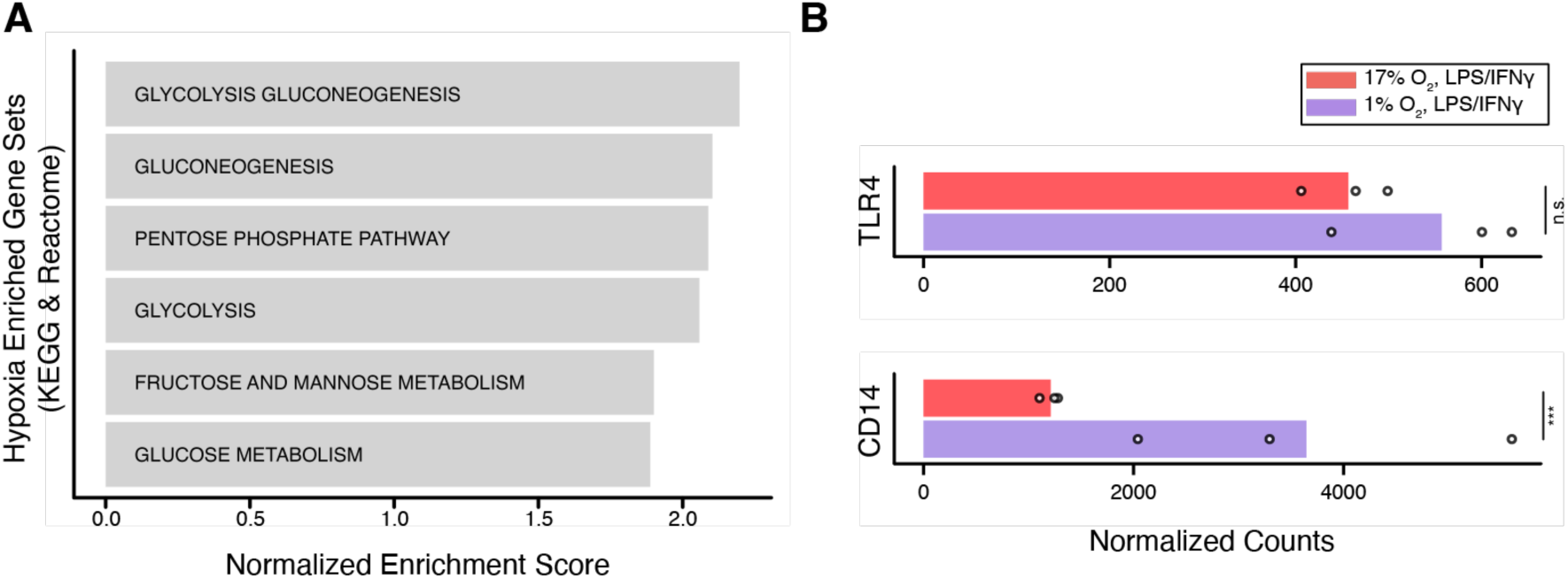
Hypoxia modifies macrophage metabolism and phagocytic specificity. A) Results from GSEA on upregulated genes in hypoxia. Data were compared to the Kyoto Encyclopedia of Genes and Genomes (KEGG) and Reactome canonical pathways gene sets. All gene sets with FDR < 0.25% are shown. B) A) Normalized read counts for TLR4 and CD14, receptors of Gram negative bacteria where points are individual replicates and bars represent the mean of those replicates. FDR value thresholds given as: *** < 0.001, and n.s. as not significant with FDR > 0.05.

**Figure S5, related to Figure 5.**
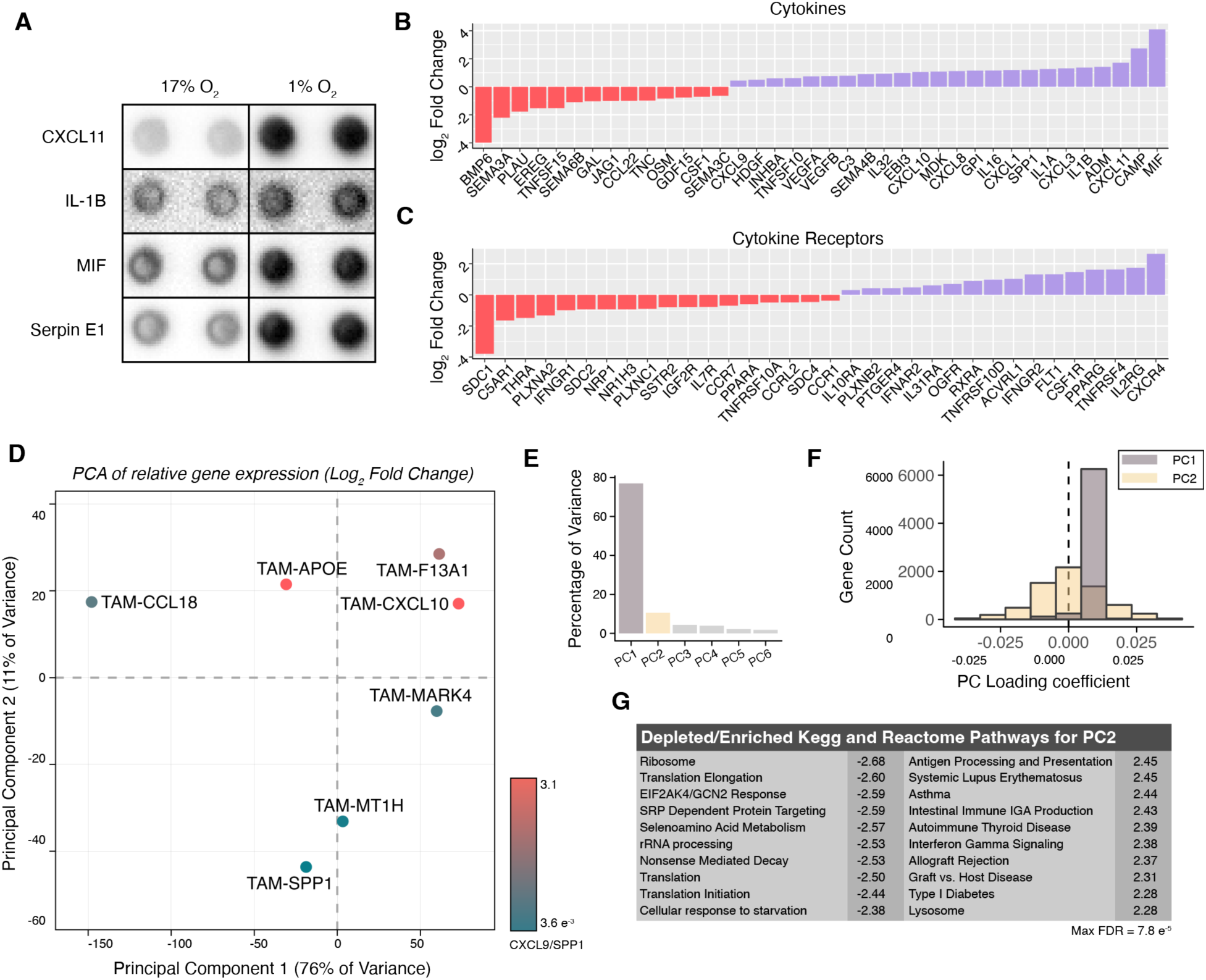
Hypoxic macrophages alter their cytokine profiles and are more correlated with tumorigenic macrophages from tumor biopsies. A) Supernatant cytokine assay comparing activated macrophages in normoxia and hypoxia. B) Differential expression values for cytokines in the comparison between activated macrophages in normoxia and hypoxia, threshold at FDR < 0.05. C) Differential expression values for cytokine receptors in the comparison between activated macrophages in normoxia and hypoxia, threshold at FDR < 0.05. D) Principal component analysis on the log_2_(fold-change) relative expression of macrophage subpopulations from Bill et al. The color of points is given as the CS ratio in the color bar provided. E) Percentage of variance for each of the 6 principal components computed for macrophage subpopulations. F) Histogram of gene loadings for the first two principal components of macrophage subpopulations. G) GSEA of gene level contributions to principal component 2. Given values are the normalized enrichment scores where negative values are for depleted pathways and positive values are for enriched pathways.

**Supplementary Data Table 1. Gene Expression and Kinetics data from macrophages.** Table contains the log_2_(fold-change) values for all genes in each of the three conditions made in this study.

